# Construction and validation of EGFP-expressing *Staphylococcus aureus* clinical strains for adhesion and internalization assays on epithelial cells

**DOI:** 10.1101/584086

**Authors:** Sana Charaoui-Boukerzaza, M. Fedy Morgene, Josselin Rigaill, Estelle Audoux, Zhiguo He, Florence Grattard, Anne Carricajo, Bruno Pozzetto, Philippe Berthelot, Elisabeth Botelho-Nevers, Paul O. Verhoeven

**Author notes:** These authors contributed equally for this work (Sana Charaoui-Boukerzaza, M. Fedy Morgene). **Corresponding author:** Paul Verhoeven (MD, PhD), Laboratory of Infectious Agents and Hygiene, University Hospital of St-Etienne, 42055 Saint Etienne Cedex 02, France.

## Abstract

**Background:** *Staphylococcus aureus* is both a major pathogen and a commensal bacterium in humans. It is able to adhere at the surface of epithelial cells of the anterior nares and can trigger its internalization inside these non-professional phagocytic cells. To better understand the interactions of clinical isolates with keratinocytes in the anterior nares, we developed and validated a one-step protocol expressing enhanced green fluorescent protein (EGFP) in *S. aureus* clinical strains with the aim to study adhesion to and internalization into mammalian cells.

**Methods:** Twenty *S. aureus* clinical isolates belonging to clonal complexes 5, 8, 30, 45, 398 were selected for one-step transformation protocol with the EGFP-encoding plasmid pBSU101. EGFP expression was analysed by flow cytometry and confocal microscopy. Wild type and isogenic EGFP-expressing strains were compared for adhesion and internalization levels by using the HaCaT cell model.

**Results:** Transformation was achieved in all the *S. aureus* strains regardless of their genetic background. The flow cytometry analysis showed that the mean proportion of EGFP-expressing bacteria was 97.2% (± 2.1) after 4h of incubation. Adhesion and internalization levels were similar in wild-type and isogenic EGFP-expressing *S. aureus* strains. Confocal laser scanning microscopy confirmed that EGFP-expressing *S. aureus* bacteria could be easily identified inside HaCaT keratinocytes.

**Conclusion:** This study reports an efficient protocol for expressing EGFP in *S. aureus* clinical strains and demonstrates that these EGFP-expressing strains are suitable for adhesion and internalization assays using HaCaT cells, which allows to perform static and dynamic *in vitro* studies of *S. aureus* colonization.

## INTRODUCTION

*Staphylococcus aureus* acts as both a leading cause of infections and a commensal bacterium in humans [1,2]. Approximately one third of the worldwide population is colonized by *S. aureus*; the anterior nares are considered as the main reservoir [1]. For long considered as an exclusive extracellular bacterium, *S. aureus* has been shown to be able to invade many non-professional phagocytic cells (NPPCs) *in vitro* but also *in vivo* during colonization or infection [3–6]. The tripartite interaction between fibronectin, the α5β1 integrin exposed at the host cell plasma membrane and fibronectin-binding proteins expressed by *S. aureus* is widely acknowledged as the main internalization pathway of *S. aureus* in NPPCs [7]. However, several others *S. aureus* virulence factors such as staphylococcal autolysin (Atl) and extracellular adherence protein (EAP) have been found to foster *S. aureus* internalization in NPPCs [8,9]. By contrast to the important knowledge generated from “laboratory” strains such as Newman or the 8325-4 derivative strains, little is known about the mechanisms driving the internalization rate and the intracellular persistence of *S. aureus* clinical isolates. Up to now, genetic manipulations in the *Staphylococcus* species relied on labour intensive protocols and only a limited number of *S. aureus* laboratory strains have been genetically modified successfully [10]. In fact, wild type *S. aureus* strains have an impenetrable restriction barrier preventing the uptake of “foreign” DNA. Monk *et al*. demonstrated that DNA plasmid isolated from *E. coli* DH10B *Δdcm* (called DC10B) bypasses the conserved type IV restriction system *SauSI*, which specifically recognizes cytosine methylated DNA, enabling transformation of *S. aureus* clinical strains [11]. This major finding has extended the possibility of genetic manipulations in *S. aureus* especially regarding clinical strains that are protected from foreign DNA.

*In vitro* cell models associated with fluorescent labelling are valuable tools to study the molecular mechanisms involved in host-pathogen interactions. However, after labelling live bacteria with fluorescent reporter, the fluorescence intensity decreases during time according to the multiplication of live bacteria (*i.e*. the quantity of fluorescent reporter is divided by two at each bacterial multiplication). The transformation of bacterial strains bearing a gene coding for a fluorescent reporter such as enhanced green fluorescent protein (EGFP) helps to overcome this limitation [12] and such bacteria could be tracked inside the cells for a few hours [13].

The aim of the present work was to develop and validate a one-step transformation protocol for constructing EGFP-expressing *S. aureus* clinical strains belonging to different genetic backgrounds without interfering with their adhesion and internalization capacities on a keratinocyte model.

## MATERIALS AND METHODS

### Microbiology procedures

The bacterial strains used in this study are listed in **Table 1**. *E. coli* (NM522 and DC10B) bacteria were grown in Luria-Bertani broth (LB) (ThermoFisher Scientific). *S. aureus* strains were growth in brain heart infusion (BHI) (Becton Dickinson) or onto tryptic soy agar (TSA) (Becton Dickinson). Culture media were incubated at 37°C with shaking at 250 rpm in case of liquid medium. When required, spectinomycin (S4014, Sigma-Aldrich) was added to the culture medium at the concentration of 120 and 125 mg/l for *S. aureus* and *E. coli*, respectively. Antimicrobial susceptibility testing of *S. aureus* clinical strains was done with the Vitek2 system (AST-P559, bioMérieux). The minimal inhibitory concentration (MIC) to spectinomycin was determined by using E-test (529240, bioMérieux). *S. aureus* strains were genotyped using spa-typing and DNA microarray (*S. aureus* genotyping kit v2.0, Alere) as previously described [14].

**Table 1.**
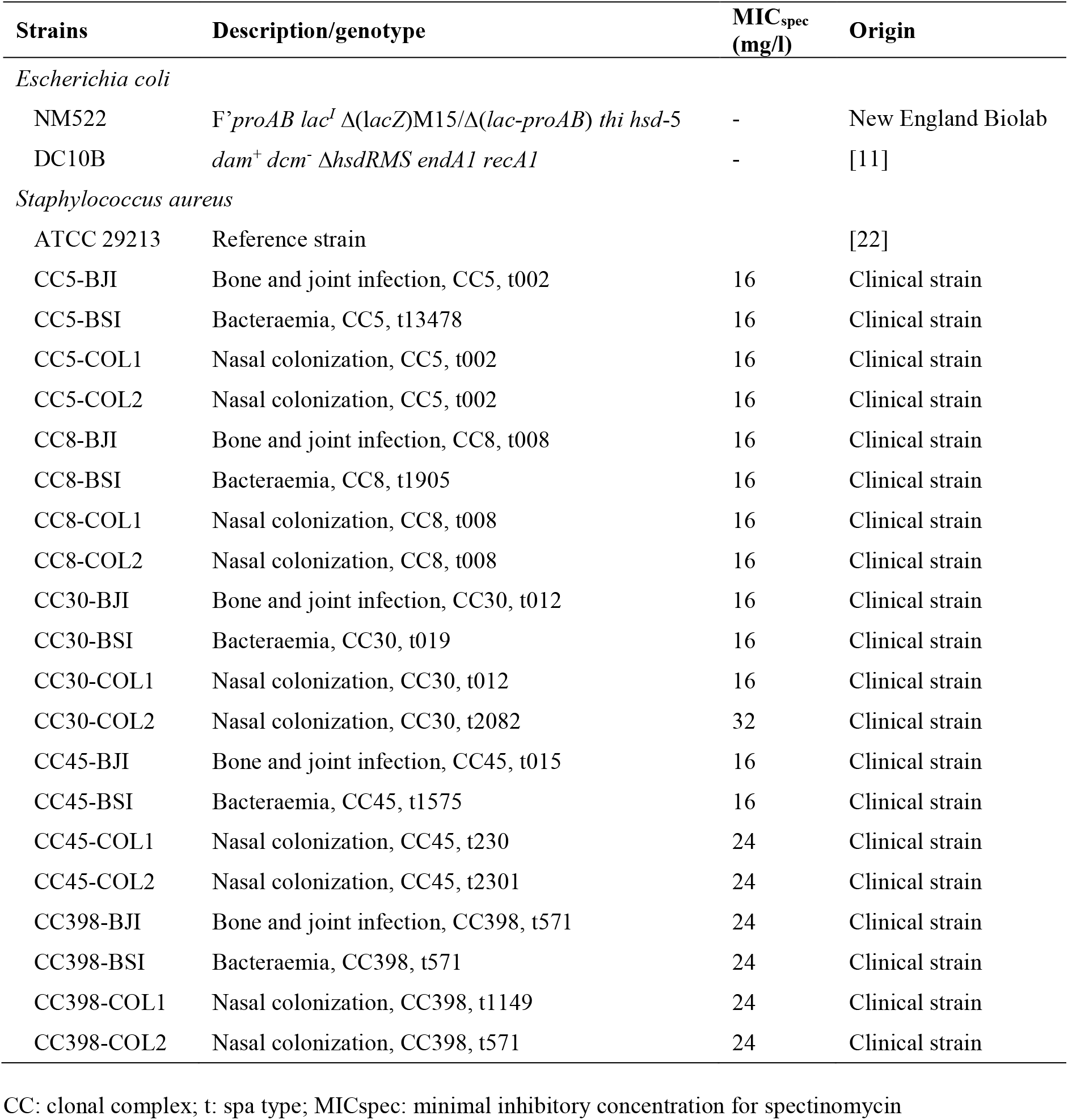
Bacterial strains used in this study.

### EGFP-encoding plasmid preparation

The pBSU101 plasmid that was kindly provided by Barbara Spellerberg was used to express EGFP in *S. aureus*. This shuttle plasmid harbours a copy of the *egfp* gene under the control of the promoter of the CAMP-factor gene (*cfb*) of *Streptococcus agalactiae*, which enables a high-level EGFP expression in several Gram-positive bacteria including *S. aureus* [12]. The pBSU101 was isolated from *E. coli* NM522 (laboratory collection) using mini spin columns (Nucleospin 740588, Macherey Nagel) according to the manufacturer’s instructions and transformed into *E. coli* DC10B, a gift from Ian Monk, by thermal shock as described [11].

### One-step transformation protocol

*S. aureus* clinical strains were inoculated in tryptic soy broth (TSB) for 24h. Overnight culture of each strain was diluted to a 0.1 optical density (OD) at 600 nm in pre-warmed TSB and incubated at 37°C with 250 rpm shaking until OD reached 0.5. Cultures were transferred to centrifuge tubes and chilled on ice for 10 minutes. The cells were harvested by centrifuging at 4500g at 4°C for 20 minutes, washed twice in ice-cold sterile water (v/v) and pelleted at 4°C. The cells were then washed in 1/10, 1/25 and 1/200 volume of ice-cold 10% sterile glycerol. Aliquots of 70 μl were stored at −80°C. For electroporation, the cells were thawed at room temperature for 10 min, centrifuged at 5000g for 3 min and re-suspended in 50 μl of sterile 10% glycerol with 500 mM sucrose. Plasmid DNA (1 μg) of pBSU101 isolated from *E. coli* DC10B was added to the cells in a sterile 0.1 cm electroporation cuvette. The cells were pulsed once at 1.8 kV and 2.5 msec time constant (MicroPulser, Biorad), outgrown in 1 ml of TSB containing 500 mM sucrose for 90 min at 37°C, spread on TSA plates containing 120 mg/l spectinomycin and incubated overnight at 37°C.

### FACS analysis of EGFP-expressing *S. aureus* clinical strains

Fluorescence stability was checked for all the transformed strains by analyzing the fluorescence means and the total fluorescent populations during the exponential phase of growth (4 hours) in BHI (Becton Dickinson) in presence of spectinomycin. A 10 μl-volume of the bacterial suspension was diluted in 490 μl of sterile distilled water and assayed by flow analysis cell sorting (FACS) (FACSCantoII, Becton Dickinson). Relative fluorescence values were determined by analyzing 50000 events for each sample using FACSDiva software (V 6.1.2, Becton Dickinson).

### Adhesion and internalization assays

Adhesion and internalization of *S. aureus* strains were studied in a model of HaCaT cells (Cell line service) that mimics nasal colonization as described previously (Rigaill 2018). Briefly HaCaT cells were seeded into 24-well plates at 1.5 x 10^5^ cells per well and maintained with serum-free medium constituted of 50% RPMI-1640 (R7388, Sigma-Aldrich) and 50% DMEM (D6429, Sigma-Aldrich) supplemented with 2% Ultroser G (Pall), 1 mM L-Glutamine (G7513, Sigma-Aldrich), 1x MEM non-essential amino acids (M7145, Sigma-Aldrich) and 1x antibiotic/antimycotic solution (A5955, Sigma-Aldrich) for three days at 37°C under 5% CO2. Cells were washed with PBS (806552, Sigma-Aldrich) and the medium was changed with fresh RPMI-1640 (R8755, Sigma-Aldrich) 24h prior to *S. aureus* inoculation. Spectinomycin (240 mg/l) was added to the wells designed to be infected with EGFP-expressing strains. Confluent cells (10^6^ cells/well) were inoculated with *S. aureus* strains in exponential phase of growth with a multiplicity of infection (MOI) of 1, and incubated for 2 hours at 37°C under 5% CO2. To quantify both adhered and internalized bacteria, cells were washed 3 times with PBS to remove free bacteria. To quantify only intracellular bacteria, the remaining extracellular bacteria were killed by adding 10 mg/l lysostaphin (Ambi Products LLC, NY, USA) for 1h at 37°C under 5% CO2. The efficacy of the lysostaphin treatment was systematically tested by plating the cell supernatant. The infected HaCaT cells were finally lysed by osmotic shock using sterile water, trypsin-EDTA (Sigma-Aldrich) and 1% Triton X-100 (Sigma-Aldrich) (2:1:1). The count of viable bacteria was carried out by plating serial dilutions of the lysates on blood agar (43041, bioMérieux) using an automatic seeder (EasySpiral Dilute, Interscience). The calculation of the bacterial loads was performed after incubation at 37°C for 24 h, using a plate-reader (Scan1200, Interscience).

### Confocal laser scanning microscopy (CLSM)

HaCaT cells were grown on glass-bottom 4-chamber culture slides (C354104, Falcon). Internalization assay was conducted as described above. After treatment with lysostaphin, cells were washed three times with PBS and fixed with 4% paraformaldehyde (Sigma-Aldrich). Cells were then permeabilized with 0.1% Triton X-100 and stained with TO-PRO-3 (T3605, Thermo Fisher Scientific) and rhodamine phalloidin (Molecular Probes). Image stacks were acquired by FluoView FV1200 CLSM equipped with UPlanSApo 60x/1.35 Oil [infinity]/0.17 FN26.5 objective and FV10-ASW4.1 software (Olympus).

### Statistical analysis

For each strain, adhesion and internalization assays were performed at least three times in two independent experiences (n = 6). The unpaired t-test was used to compare the levels of adhesion and internalization of wild-type (WT) and EGFP-expressing strains. *P* values under 0.05 were considered statistically significant. Statistical tests were performed using GraphPad Prism (v7).

## RESULTS

### One-step transformation of *S. aureus* clinical isolates

A collection of 20 *S. aureus* clinical strains isolated from bloodstream infection, prosthetic joint infection and nasal carriage, and belonging to five different clonal complexes (CC5, CC8, CC30, CC45 and CC398) were randomly selected for evaluating the transformation protocol. All the strains were susceptible to spectinomycin with MIC ranging from 16 to 48 mg/l (**Table 1**). Regardless of the genetic background or the clinical origin, all the *S. aureus* clinical strains and the reference strain ATCC 29213 were successfully transformed by electroporation with pBSU101. A single colony of each EGFP-expressing *S. aureus* strain was phenotypically selected on selective agar under UV light. The expression of EGFP was confirmed on selective agar plate under UV light and by CLSM (**Figure 1**). The constructs were all resistant to spectinomycin with a MIC of more than 128 mg/l (data not shown).

**Figure 1.**
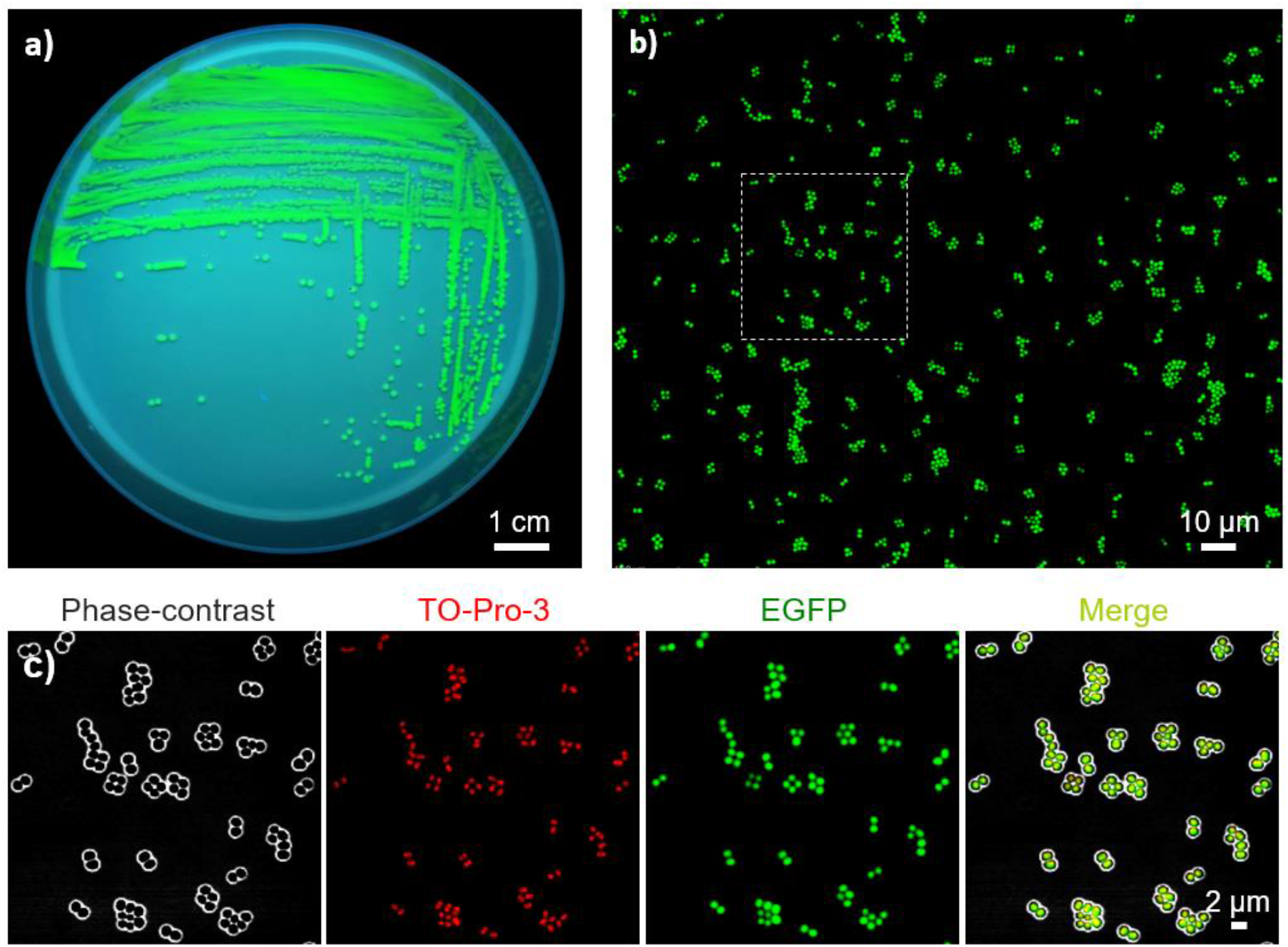
Imaging of EGFP-expressing *S. aureus* ATCC 29213 transformed with pBSU101. a) Bacterial colonies observed under UV light after 24 of incubation on TSA supplemented with 120 mg/l spectinomycin. b) Confocal laser scanning microscopy of bacterial smear on glass slide. c) Zoom magnification of bacteria observed in the dotted line box show in 1b; bacteria are depicted in phase-contrast, in the red channel (counter-staining with TO-Pro-3) and in the green channel (EGFP).

### EGFP expression by laboratory and clinical strains of *S. aureus*

EGFP expression in *S. aureus* clinical strains was assessed by FACS using the FITC channel. Figure 2a depicts the FACS diagrams of *S. aureus* ATCC 29213 transformed with pBSU101. The mean proportion of EGFP-expressing transformed bacteria was 97.2% (± 2.1) for the 20 clinical strains (**Figure 2b**). Regarding the clonal complex, the mean proportions of EGFP-expressing bacteria (± SD) was 97.9% (± 0.7), 97.9% (± 0.4), 96.8% (± 3.0), 98.0% (± 1.2) and 95.3% (± 3.0) for CC5, CC8, CC30, CC45 and CC398 strains, respectively. The mean fluorescence intensity (MFI) varied from one strain to another independently of its clonal complex or clinical origin (**Figure 2b**).

**Figure 2.**
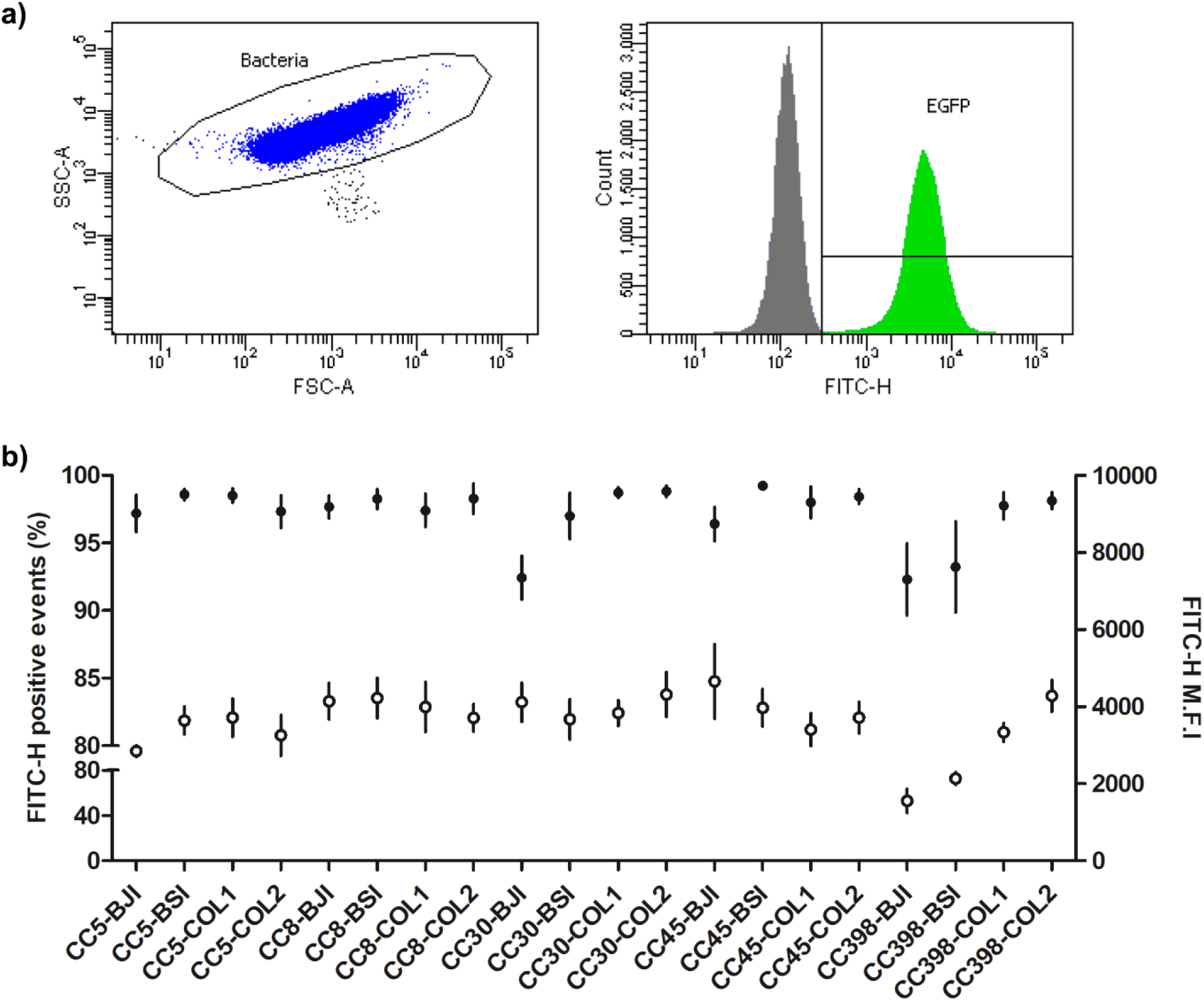
FACS analysis of EGFP-expressing *Staphylococcus aureus* strains transformed with pBSU101. a) Dot plot and FITC-H histogram of *S. aureus* ATCC 29213 transformed with pBSU101. b) FITC-H values of the 20 *S. aureus* clinical strains transformed with pBSU101. The rate of fluorescent bacteria (closed circles) and the mean fluorescence intensity (open circles) were calculated using EGFP-expressing *S. aureus* strains grown for 4h in brain heart infusion with 120 mg/l spectinomycin. Data represent the mean of 4 independent experiments for which 50000 events were recorded. Vertical bars represent the SEM. CC: clonal complex. BJI: bone and joint infection. BSI: blood stream infection. COL: nasal colonization.

### Adhesion and internalization of EGFP-expressing *S. aureus* strains

The adhesion and internalization levels of both EGFP-expressing and WT strains were assessed using a cell model of HaCaT keratinocytes inoculated at a MOI of 1. The mean ± SEM adhesion level was 6.50 ± 0.06 and 6.44 ± 0.05 log CFU/10^6^ cells for EGFP-expressing *S. aureus* and parental WT strains, respectively (**Figure 3a**). The mean ± SEM internalization level at 2h post-inoculation was 4.04 ± 0.08 and 4.03 ± 0.07 log CFU/10^6^ cells for EGFP-expressing *S. aureus* and parental WT strains, respectively (**Figure 3b**). CLSM was used to visualize EGFP-expressing strains inside HaCaT keratinocytes 2h post-inoculation: similar patterns of intracellular bacteria were observed with all the tested strains (**Figure 4**).

**Figure 3.**
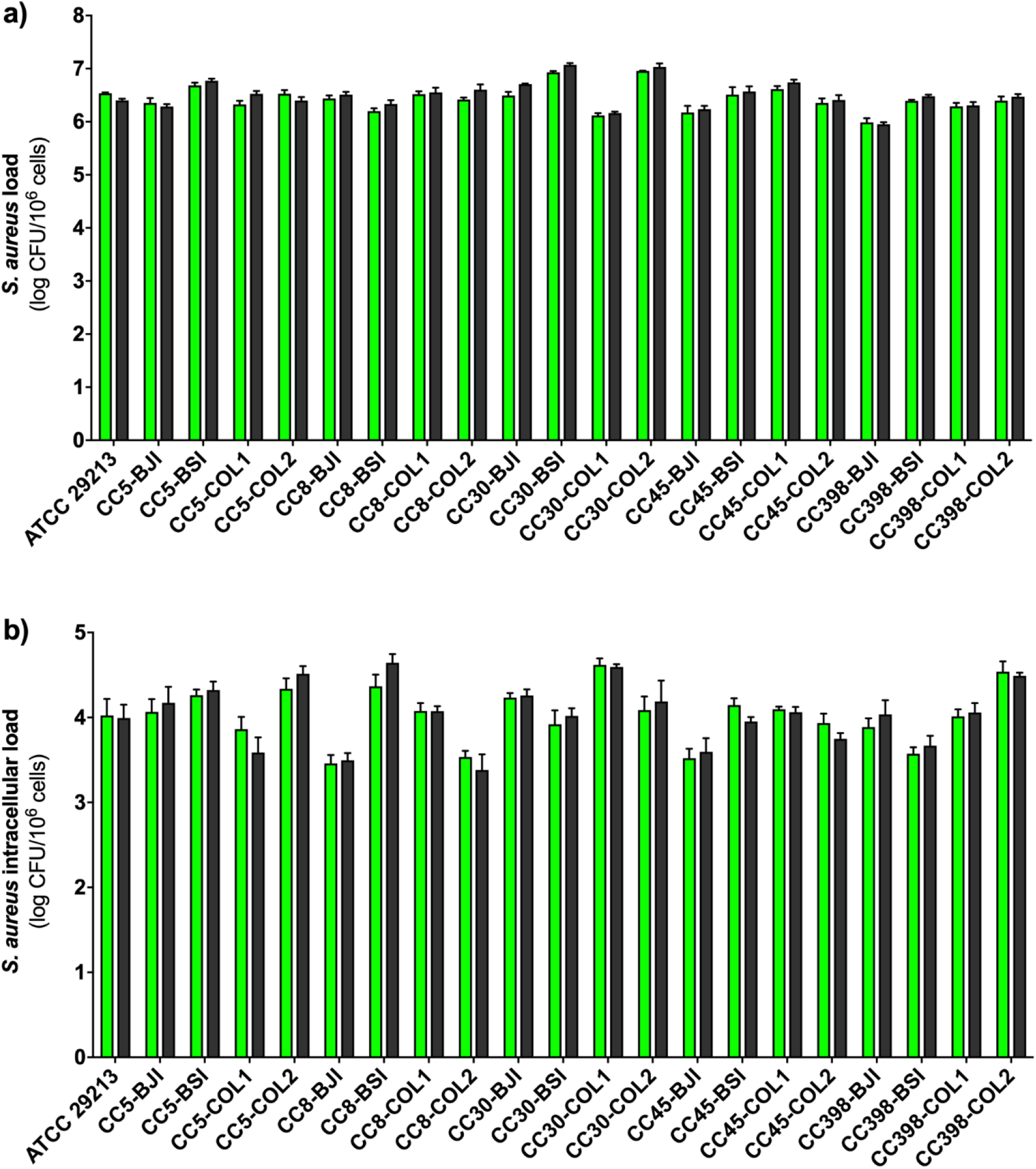
Adhesion (a) and internalization (b) levels of EGFP-expressing *S. aureus* clinical strains (green) and their parental wild-type strain (black) by using confluent monolayer of HaCaT cells inoculated with a MOI of 1. Data represent the mean of 2 independent experiments in triplicate (n = 6). Error bars correspond to the SEM. No significant difference was observed between each couple of strains (*p* > 0.05).

**Figure 4.**
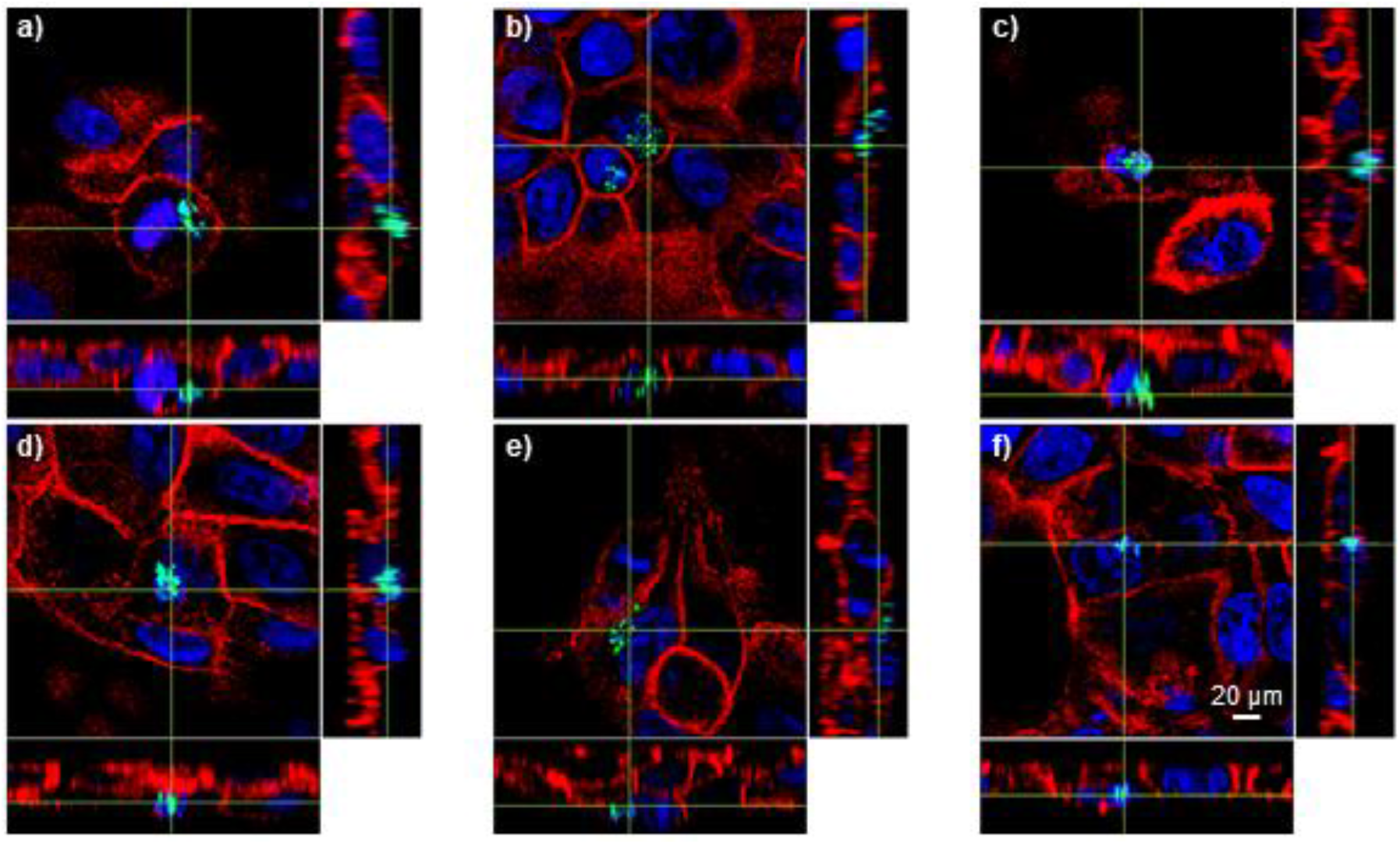
Confocal laser scanning microscopy of internalized EGFP-expressing *S. aureus* clinical strains in HaCaT cells. *S. aureus* strains CC45-COL2 (a), CC8-BJI (b), CC30-BJI (c), CC45-COL1 (d), CC398-BSI (e) and ATCC 29213 (f) are depicted in green. F-Actin and nuclei of HaCaT cells were stained by rhodamine phalloidin (red) and TO-Pro-3 (blue), respectively.

## DISCUSSION

We report herein EGFP-expression of a panel of *S. aureus* clinical strains belonging to five different genetic backgrounds by using a one-step transformation protocol based on the pBSU101 plasmid. This one-step protocol, previously described by Monk *et al*. [11] and based on the use of the *E. coli* DC10B strain, accelerates the transformation process of *S. aureus*, allowing its application to clinical isolates from different clonal complexes. For the first time, we demonstrate that the expression of EGFP decreases neither the adhesion of transformed *S. aureus* to keratinocytes nor their internalization into these cells.

The pBSU101 plasmid used in this study harbours a copy of the green fluorescent variant gene *egfp* under the control of the CAMP-factor gene (*cfb*) promoter of *Streptococcus agalactiae* [12]. By using this plasmid, the fluorescent reporter is not present at the surface of the bacteria, which avoids masking *S. aureus* surface proteins that could be involved in adhesion or internalization process. This labelling process is therefore interesting for studying interactions between *S. aureus* and host cells. By contrast, fluorescent labelling methods using dye (e.g. Vancomycin Bodipy FL) or antibodies that interact with the bacterial cell wall could modify interactions with cell host receptors [15]. In this study, we show that EGFP expressed inside the bacteria does not modify adhesion and internalization capacities of *S. aureus*. Thus we could hypothesize that the expression of the fibronectin binding proteins, the major invasion factor in *S. aureus*, is not impacted by the labelling process.

In addition, the EGFP expression enables live imaging of *S. aureus* clinical isolates because the EGFP expression driven by pBSU101 does not alter the bacterial viability [12] whereas vancomycin-based labelling has been found to exhibit a bacteriostatic activity that limits its use over the time [15]. By contrast to staining methods using peptidoglycan-binding or DNA-binding dyes, plasmid-encoded fluorescent reporters are highly specific since they are expressed only in transformed strains. Recently, Kato *et al*. [16] developed fluorescent protein tracing vectors for multicolour imaging of three clinical isolates of *S. aureus* (N315, MW2 and TY34). Furthermore, imaging of EGFP-expressing *S. aureus* avoids traditional immunostaining steps including fixation, permeabilisation and antibody labelling. Our results show that intracellular EGFP-expressing *S. aureus* clinical isolates can be easily localized by CLSM, allowing to measure the rate of *S. aureus*-invaded cells. This information is interesting since invasion rates obtained from culture experiments are calculated as mean of viable bacteria divided by total number of cells and therefore does not give any information about the distribution of *S. aureus* in the cells monolayer.

However, the use of pBSU101 for transforming clinical isolates of *S. aureus* has some limits. The EGFP fluorescence emission is known to be optimal at pH 7 but decreases almost to zero at pH 5 [17]. Since internalized *S. aureus* can be found inside phagolysosomes, the detection of these cocci can therefore be impaired. Other fluorescent proteins (e.g. red fluorescent protein) have been found to resist to pH reduction and could be useful to overcome this limitation. Besides, EGFP-encoding pBSU101 requires antibiotic selection to foster the transcription of both acetyltransferase and EGFP. Although the use of an antimicrobial agent is easy in cell models, it is not suitable in animal models. This limitation can be overpassed by chromosomic integration that ensures constitutive expression of the fluorescent reporter without the need for antibiotic pressure [18]. However, the protocol is more labour-intensive and may be difficult to apply to a great number of clinical isolates. We also observed that fluorescence intensity varied from one strain to another. Automatic quantification of bacteria based on fluorescence intensity would require a calibration step for each strain. The ability of some strains to form clump could also have an impact for automated quantification system. Nevertheless, the use of EGFP-expressing *S. aureus* strains may be easier, more specific and more accurate than live labelling of bacteria with fluorescent dyes or cell-permeable nuclear probes [15,19]. Although only EGFP was tested in this study, the pBSU101 plasmid gives opportunity to replace the *egfp* gene by another gene coding for other fluorescent proteins such as yellow fluorescent protein or mCherry that have been found suitable to label different *S. aureus* laboratory strains (RN 4220, SH1000 and SH1001) [20,21].

To the best of our knowledge, this is the first report describing a large collection of EGFP-expressing clinical strains of *S. aureus* that could represent a useful tool for studying adhesion and internalization within NPPCs such as keratinocytes. This one-step protocol could be further used to develop similar valuable tools aimed to study lifestyle of *S. aureus* during intracellular persistence or biofilm formation.

## ACKNOWLEDGMENTS

*Escherichia coli* DC10B was kindly provided by Ian Monk (Moyne Institute of Preventive Medicine, Department of Microbiology, School of Genetics and Microbiology, Trinity College Dublin, Dublin, Ireland). The pBSU101 plasmid was a gift from Prof. Barbara Spellerberg (Institute of Medical Microbiology and Hygiene, University of Ulm, Ulm, Germany).

## CONTRIBUTIONS

POV, PB and BP designed the study. SCB performed experiments for transforming *S. aureus* strains. MFM and JR performed adhesion and internalization assays. SCB, MFM and JR performed cytometry assays. HZ, JR and EA performed microscopy imaging. POV, MFM, SCB, AC and EBN analysed the results. POV, MFM, SCB, EBN, FG and BP wrote the manuscript.

